# Workload-induced changes to cell state contribute to β-cell failure in diabetes

**DOI:** 10.64898/2026.05.13.725004

**Authors:** Somesh Sai, Fenfen Liu, Austin R. Harrington, Han Zhu, Ibrahim Omar, Chun Zeng, Medhavi Mallick, Yinghui Sui, Maike Sander, Matthew Wortham

## Abstract

Insufficient insulin secretion relative to insulin demand is a key feature of type 2 diabetes (T2D). While the defects of insulin-producing β-cells in T2D are well defined, little is known about how β-cells progress from the functionally normal state to the decompensated state during the natural history of this disease. Here, we provide evidence that workload-induced β-cell overstimulation precipitates β-cell failure in T2D. We employ scRNA-seq to define workload-induced changes to β-cell transcriptional states, identifying a novel compensating state that is distinct from the stressed state of decompensated β-cells. We demonstrate a key role for the chromatin-modifying enzyme Lysine-specific demethylase 1 (Lsd1) in restraining workload-induced β-cell state transitions, indicating epigenomic control of β-cell state. Experimental manipulations that promote the compensating state accelerate β-cell failure in mouse models of diabetes. Altogether, these findings show that the compensatory response of the β-cell to increased workload becomes maladaptive over time and contributes to the pathogenesis of T2D.

## Introduction

Type 2 diabetes (T2D) is defined as hyperglycemia resulting from the inability of pancreatic β-cells to secrete enough insulin to meet the necessary amount for glucose clearance, usually in the context of insulin resistance. T2D is a complex disease characterized by genetic predisposition to β-cell dysfunction and insulin resistance that manifests as β-cell failure in response to potentiating environmental factors^1^. Recent genome-wide association studies (GWAS) of deeply phenotyped patient populations have revealed that T2D is heterogeneous due to interindividual differences in the extent that genetic background impacts β-cell function and insulin sensitivity^2–6^. Nevertheless, most individuals with T2D have insulin resistance, which exerts pressure on β-cells to increase insulin output and thereby exposes an insufficient capacity of β-cells to compensate for increased insulin demand^7,8^. While genetic background contributes to intrinsic differences in predisposition to insulin resistance and β-cell failure, the response of β-cells to environmental factors determines whether a predisposed individual will develop T2D^8^.

A mechanistic understanding of β-cell failure during the natural history of T2D will be necessary to determine how to forestall or reverse β-cell dysfunction. Extensive cross-sectional studies comparing islets from individuals with T2D or pre-T2D relative to those without diabetes have defined the major functional defects and corresponding molecular changes at these disease states^9–12^. However, there are several key limitations to cross-sectional islet studies. First, without longitudinal studies it is difficult to rule out that observed differences are intrinsic to β-cells in individuals destined to become diabetic. Furthermore, cross-sectional studies cannot avoid confounding effects of systemic hyperglycemia on β-cells^13,14^. While studies of prediabetic (HbA1c of 5.7% - 6.5%) or recently diagnosed individuals represent earlier stages of T2D pathogenesis, β-cell defects are already apparent at these time points^12,15,16^. Underscoring the challenge in defining the molecular underpinnings of β-cell dysfunction, a single cell multi-ome study of β-cell transcriptomes paired with measurements of exocytosis indicated that mRNAs correlated with insulin exocytosis in T2D were paradoxically anti-correlated with insulin exocytosis in β-cells from healthy donors^12^. Overall, very little is known about the precipitating events that cause β-cells to fail in T2D.

Insulin resistance is widely considered the initiating event in the natural history of T2D^7,8^. To preserve glucose homeostasis during insulin resistance, β-cells secrete more insulin to compensate for reduced insulin action at its target tissues^17,18^. Development of hyperglycemia does not occur unless insulin requirements outstrip the capacity of β-cells to sustain or further increase compensatory insulin secretion. The toxic effect of elevated glucose on the β-cell, termed glucotoxicity, has been well-documented and is associated with an array of cellular insults contributing to β-cell failure as described in detail elsewhere^13,16,19^. Prior to glucotoxicity, little is known about what causes β-cells to transition from the highly functional compensating state to the dysfunctional, failed state at this early stage in pathogenesis. Studies of this early disease process are hampered by a limited understanding of the fundamental mechanisms underlying β-cell compensation for insulin resistance.

In addition to the known compensatory response of the β-cell to chronic changes to insulin demand, short-term changes in nutrient state including meal feeding also evoke adaptation of the insulin secretory response. We recently focused on fasting and re-feeding of mice as a simplified model of adaptive insulin secretion, which revealed an epigenomic regulatory mechanism^20^. This program of functional adaptation corresponds to transcriptional changes in nutrient response genes that are regulated in part through acetylation of histone 3 lysine 27 (H3K27ac), an activating histone modification, at cognate regulatory elements. Briefly, feeding leads to accumulation of H3K27ac at the regulatory elements of nutrient response genes, thereby increasing expression of these genes and enhancing the insulin secretory response to subsequent stimuli. We found that feeding-induced H3K27ac deposition occurs at sites bound by the chromatin-modifying enzyme Lysine-specific demethylase 1 (Lsd1). Mechanistic studies in the fasting-refeeding model showed that Lsd1 restrains adaptive insulin secretion by dampening feeding-induced accumulation of H3K27ac^20^. Comparative studies suggested that β-cells employ shared mechanisms to adjust the strength of the insulin secretory response to short- and long-term changes in nutrient state corresponding to anticipated insulin requirements per β-cell (β-cell workload hereafter)^20^. In human islets, we found that GWAS variants associated with T2D were enriched at LSD1-bound H3K27ac peaks, suggesting there is a causative link between the LSD1-regulated adaptation program and T2D pathogenesis. However, as our prior studies focused on functional adaptations to feeding^20^, it was unclear how this epigenomic regulatory mechanism for adaptive insulin secretion would impact β-cell failure in T2D.

Here, we show that experimentally activating the Lsd1-controlled adaptation program has the paradoxical effect of accelerating β-cell failure during insulin resistance. Analysis of β-cell transcriptomes at single cell resolution in models of insulin resistance with and without β-cell *Lsd1* deletion revealed that increased workload alters β-cell state in a manner that is normally restrained by Lsd1. Longitudinal analysis and pseudotime cell ordering in models of unsustainable workload predict that an intermediate ‘compensating’ β-cell state is a precursor to the ‘stressed’ state, suggesting that the compensatory response to increased workload becomes maladaptive when it is chronically activated.

## Results

### Lsd1 inactivation accelerates β-cell failure during insulin resistance

To understand how the Lsd1-regulated adaptation program impacts β-cell compensation in insulin resistance, we deleted *Lsd1* in β-cells of juvenile *db/db* mice using *Pdx1-CreER* (*db/db Lsd1^Δβ^* mice hereafter; **Fig. 1a**, **Extended Data Fig. 1a**) then monitored diabetes progression relative to Lsd1-intact *db/db* mice (*db/db Lsd1^fl/+β^* hereafter; **Fig. 1b**, **Extended Data Fig. 1b**). To assess whether previously described *Lsd1^Δβ^* phenotypes^20^ are recapitulated at this early deletion time point, *Lsd1* was inactivated in parallel in β-cells of lean *db/+* mice. On this strain background and treatment scheme, Lsd1-intact *db/db* mice slowly and progressively developed hyperglycemia over 24 weeks (**Fig. 1b**). Diabetes onset was accelerated by β-cell *Lsd1* inactivation, with *db/db Lsd1^Δβ^* mice developing overt hyperglycemia by 10 weeks of age. Longitudinal analysis revealed exaggerated glucose intolerance in 9-week-old *db/db Lsd1^Δβ^* mice compared to Lsd1-intact *db/db* controls and little effect of acute *Lsd1* inactivation on glucose tolerance in 6- to 7-week-old *db/db* mice (**Fig. 1c**, **Extended Data Fig. 1c-f**). Impaired glucose tolerance following β-cell *Lsd1* inactivation in *db/db* mice corresponded to reduced fasting insulin levels, consistent with β-cell dysfunction (**Fig. 1d**). Worsening glucose homeostasis in *db/db Lsd1^Δβ^* mice could not be attributed to reductions in insulin sensitivity or pancreatic insulin content (**Extended Data Fig. 1g, h**). Quantification of β-cell mass revealed a modest reduction in *db/db Lsd1^Δβ^* mice compared to *db/db Lsd1^fl/+β^* mice (**Extended Data Fig. 1i**). These phenotypes in a preclinical model of obesity and insulin resistance are in stark contrast to effects of β-cell *Lsd1* inactivation in lean mice, which causes hypoglycemia (**Fig. 1b**) as previously described^20^. Therefore, Lsd1 has opposing effects on glycemia depending on metabolic context.

**Figure 1.**
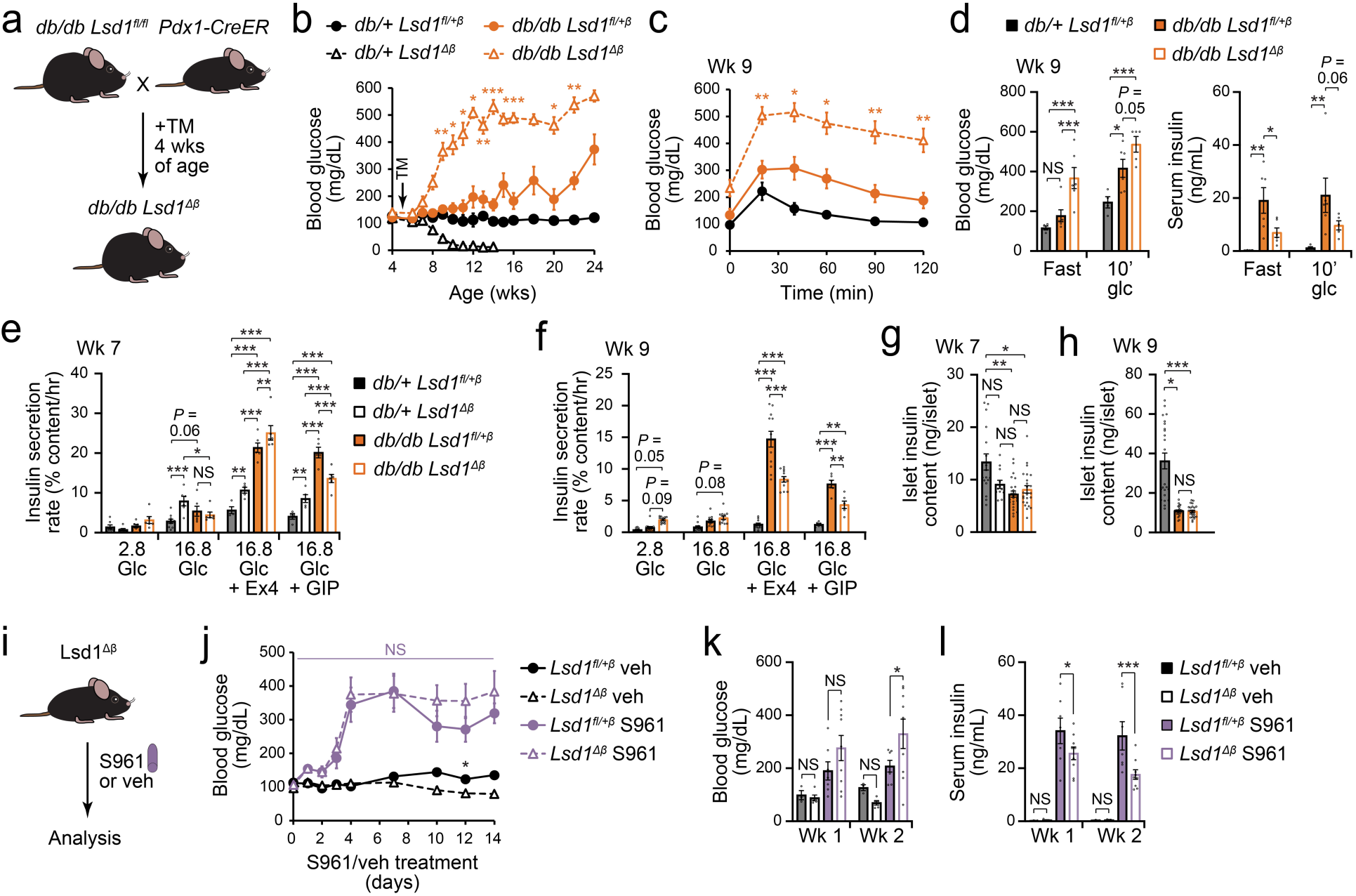
*Lsd1* inactivation impairs β-cell adaptation to insulin resistance. **(a)** Schematic of alleles and treatments used to inactivate *Lsd1* in *db/db* mice. TM, tamoxifen; wks, weeks. **(b)** Time course of ad libitum-fed blood glucose levels in TM-treated mice of the indicated genotypes. *db/+ Lsd1^fl/+β^*: *n* = 7 mice, *db/db Lsd1^fl/+β^*: *n* = 8 mice, *db/db Lsd1^Δβ^*: *n* = 11 mice, *db/+ Lsd1^Δβ^*: *n* = 14 mice. \**p*<0.05, \*\**p*<0.01, \*\*\**p*<0.001 between *db/db Lsd1^fl/+β^* and *db/db Lsd1^Δβ^* mice. **(c)** Glucose tolerance tests in TM-treated mice of the indicated genotypes. *db/+ Lsd1^fl/+β^*: *n* = 6 mice, *db/db Lsd1^Δβ^*: *n* = 11 mice, *db/db Lsd1^fl/+β^*: *n* = 13 mice. \**p*<0.05, \*\**p*<0.01 between *db/db Lsd1^fl/+β^*and *db/db Lsd1^Δβ^* mice. **(d)** Serum insulin and blood glucose (glc) levels in mice of the indicated genotypes following a 6-hour fast or 10 min following intraperitoneal injection of glucose. *db/+ Lsd1^fl/+β^*fast and 10’ glucose: *n* = 4 mice, *db/db Lsd1^Δβ^* 10’ glucose: *n* = 5 mice, all other groups: *n* = 6 mice. **(e** and **f)** Static insulin secretion assays for islets from the indicated genotypes of mice stimulated with the indicated glucose (glc) concentrations (in mM) with or without 10 nM of exendin-4 (Ex4) or GIP at 7 wks (e) and 9 wks (f) of age. *db/+ Lsd1^fl/+β^* 16.8 mM glc + Ex4 or GIP wk 7: *n* = 4 pools of 10 islets each, *db/+ Lsd1^Δβ^* 16.8 mM glc + Ex4 or GIP wk 7 and *db/db Lsd1^fl/+β^* 16.8 mM glc + Ex4 wk 7: *n* = 5 islet pools, *db/+ Lsd1^fl/+β^* 16.8 mM glc wk 7: *n* = 10 islet pools, *db/+ Lsd1^fl/+β^* 2.8 mM glc wk 9 and *db/db Lsd1^fl/+β^* 16.8 mM glc wk 9: *n* = 11 islet pools, *db/db Lsd1^fl/+β^* 2.8 mM glc wk 9, *db/db Lsd1^Δβ^*2.8 mM glc or 16.8 mM glc wk 9, *db/+ Lsd1^fl/+β^* 16.8 mM glc wk 9, and all genotypes 16.8 mM glc + Ex4 wk 9: *n* = 12 islet pools, all other groups: *n* = 6 islet pools. **(g** and **h)** Islet insulin content for islets from the indicated genotypes of mice. *db/+ Lsd1^Δβ^* wk 7: *n* = 12 pools of 10 islets each, *db/+ Lsd1^fl/+β^* wk 7: *n* = 17 islet pools, *db/+ Lsd1^Δβ^* wk 9: *n* = 23 islet pools, all other groups: *n* = 24 islet pools. **(i)** Schematic of S961 administration via transplanted minipumps (20 nmol/week). Veh, vehicle. **(j)** Time course of ad libitum-fed blood glucose levels in TM-treated *Lsd1^fl/+^*; *Pdx1-CreER* mice (*Lsd1^fl/+^*^β^) and TM-treated *Lsd1^fl/fl^*; *Pdx1-CreER* mice (*Lsd1^Δβ^*) administered S961 or vehicle. *Lsd1^fl/+^*^β^ veh: *n* = 3, *Lsd1^Δβ^* veh: *n* = 5, *Lsd1^fl/+^*^β^ S961: *n* = 7, *Lsd1^Δβ^*S961, *n* = 9. NS, not significant between S961-treated *Lsd1^fl/+^*^β^ and *Lsd1^Δβ^* mice. \**p*<0.05 between *Lsd1^fl/+^*^β^ and *Lsd1^Δβ^* mice **(k** and **l)** Blood glucose levels (k) and serum insulin levels (l) after a 6-hour fast in *Lsd1^fl/+^*^β^ and *Lsd1^Δβ^*treated with S961 or vehicle for the indicated weeks. *Lsd1^fl/+^*^β^ veh: *n* = 3, *Lsd1^Δβ^* veh: *n* = 5, *Lsd1^fl/+^*^β^ S961: *n* = 7, *Lsd1^Δβ^* S961, *n* = 9. Significance was determined by one-way ANOVA followed by Student’s t-test with Welch’s correction for unequal variance as necessary followed by Dunnett’s multiple comparisons test (g and h) or by two-way ANOVA for treatment or genotype interaction with time or stimulation condition followed by Sidak’s (b, c, j) or Benjamini, Krieger and Yekutieli multiple comparisons test (d - f, k, l). \**p*<0.05, \*\**p*<0.01, \*\*\**p*<0.001; NS, not significant.

Given the stark differences in onset of glucose intolerance and diabetes between β-cell *Lsd1*-deficient and Lsd1-intact *db/db* mice, we predicted that impaired insulin secretion could contribute to disease progression following β-cell *Lsd1* inactivation. We therefore assessed β-cell function using static insulin secretion assays prior to and coincident with effects of β-cell *Lsd1* inactivation on glucose tolerance at 7 and 9 weeks of age, respectively (**Fig. 1e, f**). Although glucose-stimulated insulin secretion was not impaired by β-cell *Lsd1* inactivation, *db/db Lsd1^Δβ^*islets exhibited basal insulin hypersecretion not seen in Lsd1-intact *db/db* mice (**Fig. 1f**). As these islet insulin secretion results contrast with circulating insulin levels, we reasoned that β-cell *Lsd1* inactivation could impair responses to non-glucose secretagogues and therefore assessed potentiation of glucose-stimulated insulin secretion by the incretin hormones GIP and GLP-1 (assessed with exendin-4). Incretin-potentiated insulin secretion was greatly enhanced in *db/db Lsd1^fl/+β^* islets compared to islets from lean *db/+ Lsd1^fl/+β^* mice; however, following *Lsd1* inactivation in *db/db* mice there was reduced GIP responsiveness and over time an attenuation of the GLP-1 response (**Fig. 1e, f**). While insulin content was reduced in islets from *db/db Lsd1^fl/+β^* mice compared to those from lean *db/+ Lsd1^fl/+β^* mice, there was no difference between *db/db Lsd1^Δβ^* and *db/db Lsd1^fl/+β^* islets (**Fig. 1g, h**). Overall, these observations suggest that insufficient potentiation of insulin secretion by GIP and GLP-1 following β-cell *Lsd1* inactivation contributes to accelerated diabetes pathogenesis in *db/db* mice, when insulin demand is very high.

Our observations that *Lsd1* inactivation has opposing effects on incretin-potentiated insulin secretion and glycemia in lean^20^ and obese states (**Fig. 1b, e, and f**) raises the question of how Lsd1 function diverges based on metabolic context. We reasoned that differences in the systemic environment could account for context-dependent functions of Lsd1. During obesity, β-cells are exposed to elevated free fatty acids^1^, chronic low-grade inflammation^21^, and increased workload due to insulin resistance^7^. To assess the requirement of increased workload for *Lsd1* inactivation to precipitate β-cell dysfunction, we employed S961, which blocks the insulin receptor to acutely cause severe insulin resistance and hyperglycemia in lean mice (**Fig. 1i, j**). Indeed, measurements of circulating insulin in S961-treated *Lsd1^Δβ^*mice revealed impaired β-cell adaptation compared to S961-treated controls (**Fig. 1k, l**). Taken together with our findings in the *db/db* model, these observations suggest that chronic stimulation of β-cells predisposes them to failure in a time-dependent manner. While increased workload can overstimulate β-cells over time, concomitant *Lsd1* deletion further stimulates β-cells (**Fig. 1f**) to potentiate the effect of workload and accelerate decompensation.

### Lsd1 dampens workload-induced changes to β-cell states

To identify molecular mechanisms whereby Lsd1 forestalls β-cell failure during insulin resistance, we performed single cell RNA-sequencing (scRNA-seq). Islets were analyzed from Lsd1-intact and β-cell *Lsd1* knockout mice treated with S961 or on the *db/db* background prior to and coincident with the onset of glucose intolerance (**Fig. 2a** and **Extended Data Fig. 1c**). Clustering of β-cells (*Ins1/2*-expressing cells) identified four β-cell states (β-1 through β-4), which we categorized based on enriched gene ontologies (GOs) and pathways for mRNAs overexpressed in each state (**Fig. 2b-d**)^22^. β-1 ‘homeostatic’ cells expressed high levels of β-cell-characteristic transporters (e.g., *Slc2a2* and *Slc30a8*) and metabolic enzyme genes (*G6pc2*), while β-4 ‘proliferating’ cells were enriched for transcripts involved in DNA replication (*Pcna* and *Top2a*) and cell cycle progression (*Ccna2, Ccnb1*, and *Cdk1*; **Fig. 2c, d**, **Extended Data Fig. 2a**, and **Supplemental Table 1**). β-2 ‘compensating’ and β-3 ‘stressed’ cells were characterized by high expression of mRNAs related to protein folding (*Sdf2l1* and *Fkbp11*), with β-3 ‘stressed’ cells additionally highly expressing genes related to cellular stress responses (*Gpx3*, *Hspa5*, and *Txnip*; **Fig. 2c, d**).

**Figure 2.**
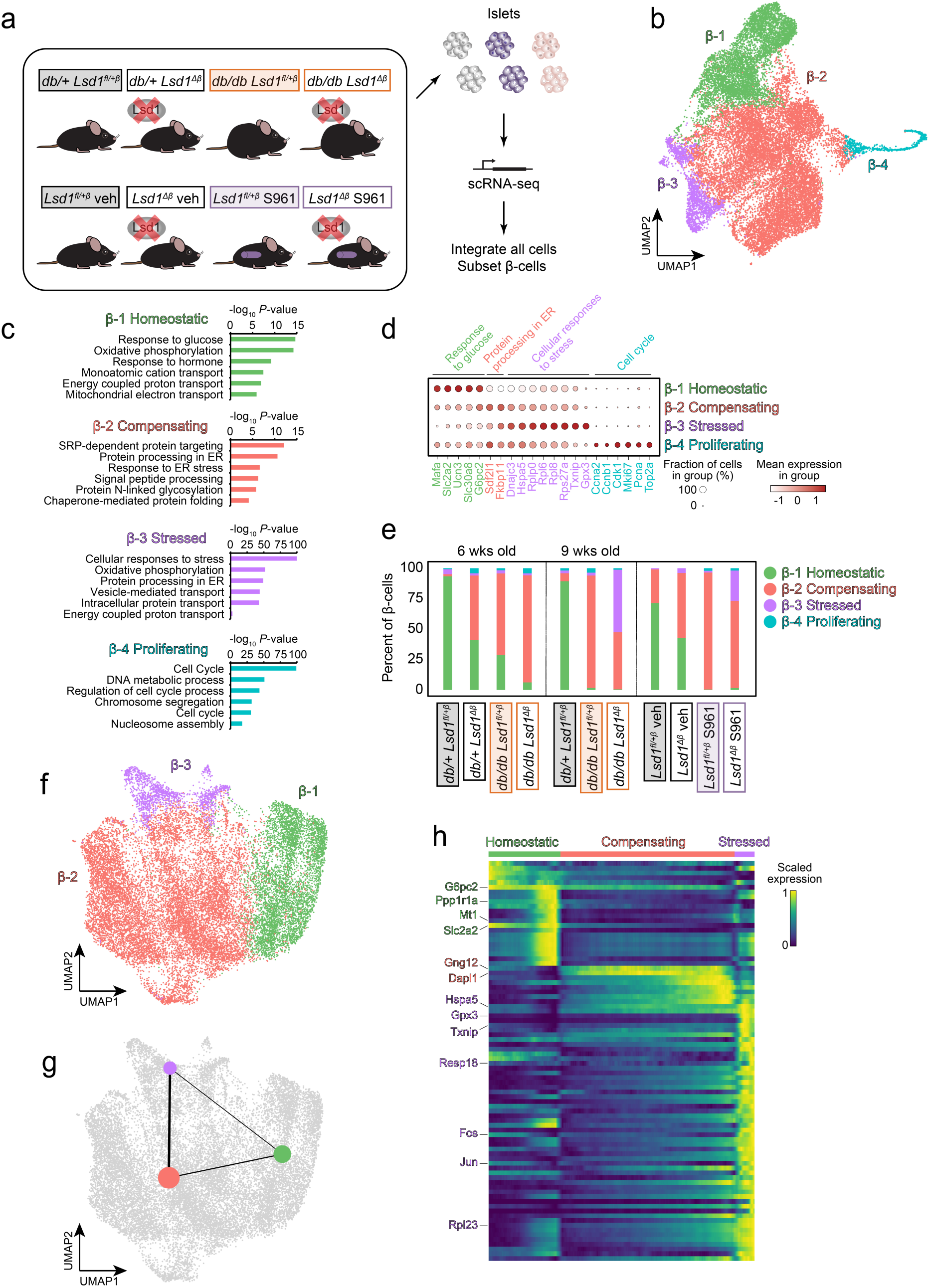
Lsd1 restrains workload-induced changes to β-cell states. **(a)** Schematic of experimental groups and workflow. Mice were TM-treated at either 4 wks of age (*db/db* experiment) or between 10-14 wks of age (S961 experiment) then islets were harvested 1 and 4 wks later (*db/db* experiment) or 2 wks later (S961 experiment). TM, tamoxifen; wks, weeks. **(b)** UMAP of clustered β-cells colored by state. UMAP, uniform manifold approximation and projection. **(c)** Enriched gene ontologies and pathways for mRNAs more highly expressed in each state relative to all other β-cells. The top six most significantly enriched categories are shown. **(d)** Dot plot showing expression of representative mRNAs from gene ontologies and pathways selected from (c) for each β-cell state. Font color indicates the state most highly expressing each mRNA. **(e)** Abundance of each β-cell state as a proportion of total β-cells for the indicated treatments and genotypes. **(f)** UMAP of clustered nonproliferating β-cells colored by state. **(g)** PAGA graph superimposed on a UMAP showing predicted paths between the β-cell states from (f). PAGA, partition-based graph abstraction. **(h)** Expression changes for selected mRNAs along the diffusion pseudotime trajectory from (g). Cells were ordered as in ^27^. Full labeling is shown in Extended Data Fig. 2c.

To determine how the metabolic environment affects β-cell states, we assessed their abundances in the different groups (**Fig. 2e** and **Extended Data Fig. 2b**). β-1 ‘homeostatic’ cells were predominant in insulin-sensitive, Lsd1-intact control mice. In insulin resistance, β-2 ‘compensating’ cells accumulated at the expense of β-1 ‘homeostatic’ cells. Analysis of *Lsd1*-deficient groups revealed context-dependent effects of Lsd1 on β-cell states. *Lsd1* deletion in lean, normoglycemic mice increased the abundance of β-2 ‘compensating’ cells and depleted β-1 ‘homeostatic’ cells akin to insulin resistance. In insulin resistant mice with preserved glucose tolerance (i.e., 6-week-old *db/db* mice; **Extended Data Fig. 1c**), inactivation of *Lsd1* caused further accumulation of β-2 ‘compensating’ cells (**Fig. 2e**). In contexts of glucose intolerance or hyperglycemia Lsd1 prevented the emergence of β-3 ‘stressed’ cells (**Fig. 2e**).

Based on the magnitude of β-cell state changes over 2-3 weeks of insulin resistance, we speculated these arose in part due to interconversion among β-1 through β-3 cells. We therefore performed a cell ordering approach using PAGA (partition-based graph abstraction) and diffusion pseudotime to predict transitions between β-cell states (**Fig. 2f-h** and **Extended Data Fig. 2c**). PAGA predicted a high-confidence path between β-2 ‘compensating’ cells and β-3 ‘stressed’ cells, suggesting that β-2 ‘compensating’ cells are in an intermediate state. Together with the above-described changes to β-cell state abundances (**Fig. 2e**), these observations indicate that Lsd1 restrains cell state transitions caused by increased β-cell workload. In this way, *Lsd1* inactivation accelerates β-cell failure in response to unsustainable workloads by potentiating the effects of insulin resistance and hyperglycemia on β-cell states.

### ‘Compensating’ β-cells accumulate in diverse models of increased workload

We next assessed the generalizability of the observed β-cell state transitions in response to increased workload. Novel datasets were generated and integrated with relevant published studies^23–24^ to build a scRNA-seq atlas comprising 124,191 islet cells, from which β-cells were re-integrated and clustered to identify and categorize β-cell states as above (**Fig. 3a-c**, **Extended Data Fig. 3a**, and **Supplemental Table 2)**. β-cells from control samples clustered together on the atlas embedding, indicating successful integration (**Extended Data Fig. 3b**). β-cells clustered into six total states, with the vast majority of β-cells belonging to one of three states: β-1 ‘homeostatic’ cells (28% of all β-cells), β-2 ‘compensating’ cells (55%), and β-3 ‘stressed-immature’ cells (15%; **Fig. 3b-d** and **Extended Data Fig. 4a-c**). GO term and pathway enrichment of mRNAs characteristic of β-1 through β-4 cells from this islet atlas were similar to those for the β-cell states from models of increased workload with and without β-cell *Lsd1* deletion (**Fig. 2c**). All β-cell states except for β-6 ‘batch-specific’ cells were detected in multiple samples, further supporting the robustness of sample integration (**Fig. 3e** and **Extended Data Fig. 4b, c**). β-1 ‘homeostatic’ cells highly expressed mRNAs related to β-cell maturity and function (e.g., *Glp1r*, *Mafa*, *Ppp1r1a*, and *Slc2a2*) while β-3 ‘stressed-immature’ cells were depleted of β-cell maturity transcripts (*Mafa* and *Ucn3*) and highly expressed genes involved in protein synthesis and processing (*Dnajc3*, *Hspa5*, *Rpl6*, and *Rpl8*) as well as stress responses (*Txnip* and *Cd81*; **Fig. 3c, d** and **Supplemental Table 3**). β-2 ‘compensating’ cells highly expressed mRNAs involved in ‘protein folding’ (*Fkbp11* and *Sdf2l1*; **Fig. 3c**) and expressed intermediate levels of transcripts characteristic of either β-1 or β-3 cells (**Fig. 3d**). Analysis of β-cell state abundances across samples showed β-3 ‘stressed-immature’ cells were most abundant in samples from chronically hyperglycemic (STZ-treated or *ob/ob*) or developmentally immature (3-week-old) mice (**Fig. 3e** and **Extended Data Fig. 4d**), suggesting these correspond to previously-described ‘dedifferentiated’ or ‘immature’ β-cells^24–26^. With the exception of chronically hyperglycemic STZ-treated mice, models of increased β-cell workload accumulated β-2 ‘compensating’ cells at the expense of β-1 ‘homeostatic’ cells relative to controls (**Fig. 3e**). In fasted and re-fed mice these abundance changes occur within hours, suggesting β-cells transition between β-1 and β-2 states based on the relative surplus or shortfall of insulin (**Fig. 3e**). The accumulation of β-2 ‘compensating’ cells and depletion of β-1 ‘homeostatic’ cells was reproduced in an independent diet-induced obesity dataset mapped to the atlas embedding through label transfer (**Extended Data Fig. 4e**). Importantly, β-cell state classifications were not driven by differences in cell quality metrics among β-cell states (**Extended Data Fig. 4a, f-h**). Overall, these data indicate that increased workload causes β-cells to transition to a ‘compensating’ state that is distinct from the failed, dedifferentiated state associated with diabetes.

**Figure 3.**
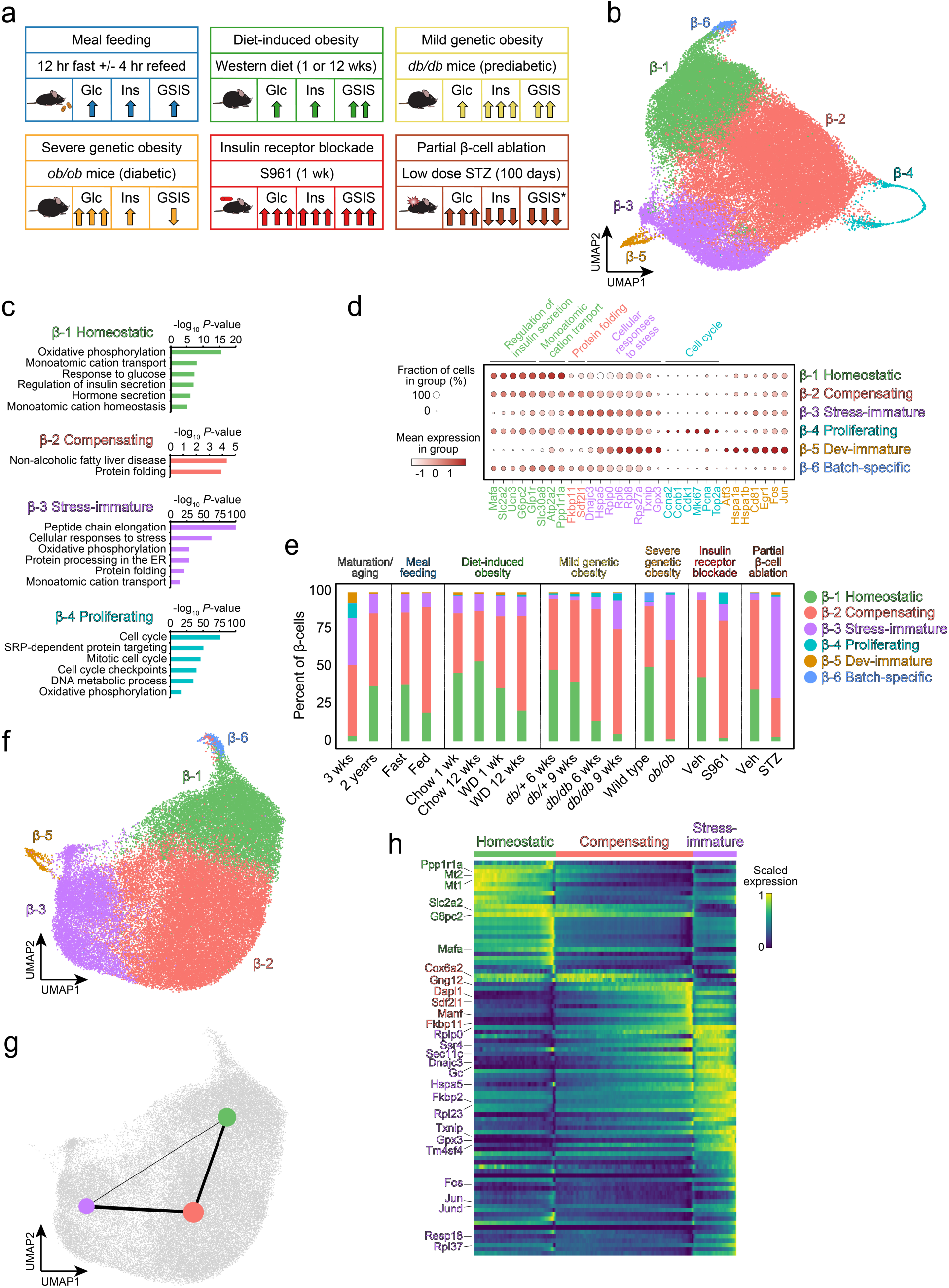
Diverse models of increased workload impact β-cell states. **(a)** Schematic of experimental groups analyzed by scRNA-seq. For each group, relative changes to circulating glucose (glc) and insulin (ins) levels, or to GSIS by isolated islets, are depicted with arrows approximating effect sizes. *inferred based on circulating insulin levels. GSIS, glucose-stimulated insulin secretion. STZ, streptozotocin. **(b)** UMAP of re-integrated and clustered β-cells colored by state. UMAP, uniform manifold approximation and projection. **(c)** Enriched gene ontologies and pathways for mRNAs more highly expressed in each state relative to all other β-cells. The top six most significantly enriched categories are shown. **(d)** Dot plot showing expression of representative mRNAs from gene ontologies and pathways selected from (c) for each β-cell state. Font color indicates the state most highly expressing each mRNA. **(e)** Abundance of each β-cell state as a proportion of total β-cells for the indicated treatments and genotypes. **(f)** UMAP of clustered nonproliferating β-cells colored by state. **(g)** PAGA graph superimposed on a UMAP showing predicted paths between the three most abundant β-cell states. PAGA, partition-based graph abstraction. **(h)** Expression changes for selected mRNAs along the diffusion pseudotime trajectory for the three most abundant β-cell states. Cells were ordered as in ^27^. Full labeling is shown in Extended Data Fig. 5.

The above observations suggest that diabetes pathogenesis involves stepwise conversion of ‘homeostatic’ β-cells first to the ‘compensating’ state then to the ‘stressed-immature’ state. To predict transitions between the β-1 ‘homeostatic’ state and workload-regulated β-2 ‘compensating’ or β-3 ‘stressed-immature’ β-cell states, we performed PAGA as above and generated cell trajectories using diffusion pseudotime (**Fig. 3f-h** and **Extended Data Fig. 5**)^27^. PAGA inferred a major path connecting β-1 ‘homeostatic’ cells and β-3 ‘stressed-immature’ cells via the β-2 ‘compensating’ state (**Fig. 3g**). Gene expression along this diffusion pseudotime trajectory showed progressive downregulation of transcripts characteristic of β-1 ‘homeostatic’ β-cells and upregulation of mRNAs enriched in β-3 ‘stressed-immature’ β-cells, suggesting the ‘compensating’ state is a transitional state between healthy and failed β-cells (**Fig. 3h** and **Extended Data Fig. 5**). Overall, these observations suggest the natural history of diabetes pathogenesis involves early transition of β-cells from the ‘homeostatic’ to the ‘compensating’ state due to increased workload, followed by later transition to a stressed state corresponding to failure of overstimulated β-cells.

### β-cell states exhibit different activities of gene regulatory networks governing β-cell identity and function

To predict transcription factors regulating β-cell state transitions, we constructed gene regulatory networks (GRNs) in the islet atlas using SCENIC^28^ (**Fig. 4a** and **Extended Data Fig. 6a**), identifying 124 β-cell regulons that clustered into 5 inter-correlated modules (**Fig. 4b**). As expected, activity of β-cell lineage-determining TFs (LDTFs) such as Neurod1, Nkx6.1, and Pdx1^29^ were closely correlated with one another and anti-correlated with TFs involved in stress-induced loss of β-cell identity such as Ascl1^30^ and Bach2^31^ (**Fig. 4b**). Visualization of TFs and their target genes as a network revealed a high degree of target gene co-regulation for these β-cell LDTFs (**Fig. 4c**), consistent with the high degree of overlap for corresponding cistromes from islet ChIP-seq data^29,32,33^. Comparison of regulon activities between β-cell states identified state-dependent activities for several TFs with known functions in the β-cell^34–37^ (**Fig. 4d** and **Extended Data Fig. 6b**). For example, the progressive reduction of Neurod1 activity in β-2 ‘compensating’ and β-3 ‘stressed-immature’ cells compared to β-1 ‘homeostatic’ cells (**Fig. 4d**) suggests Neurod1 function is disrupted early in diabetes pathogenesis. Other predicted regulators such as Kdm2b, Meox1, and Pknox1 have not been characterized in β-cells and thus are strong candidates for regulation of β-cell state in response to altered workload. Overall, by establishing the relevance of workload-regulated β-cell states to diabetes pathogenesis and cataloguing GRNs associated with state transitions, this work provides a foundation for future mechanistic studies of β-cell adaptation and failure over the natural history of T2D.

**Figure 4.**
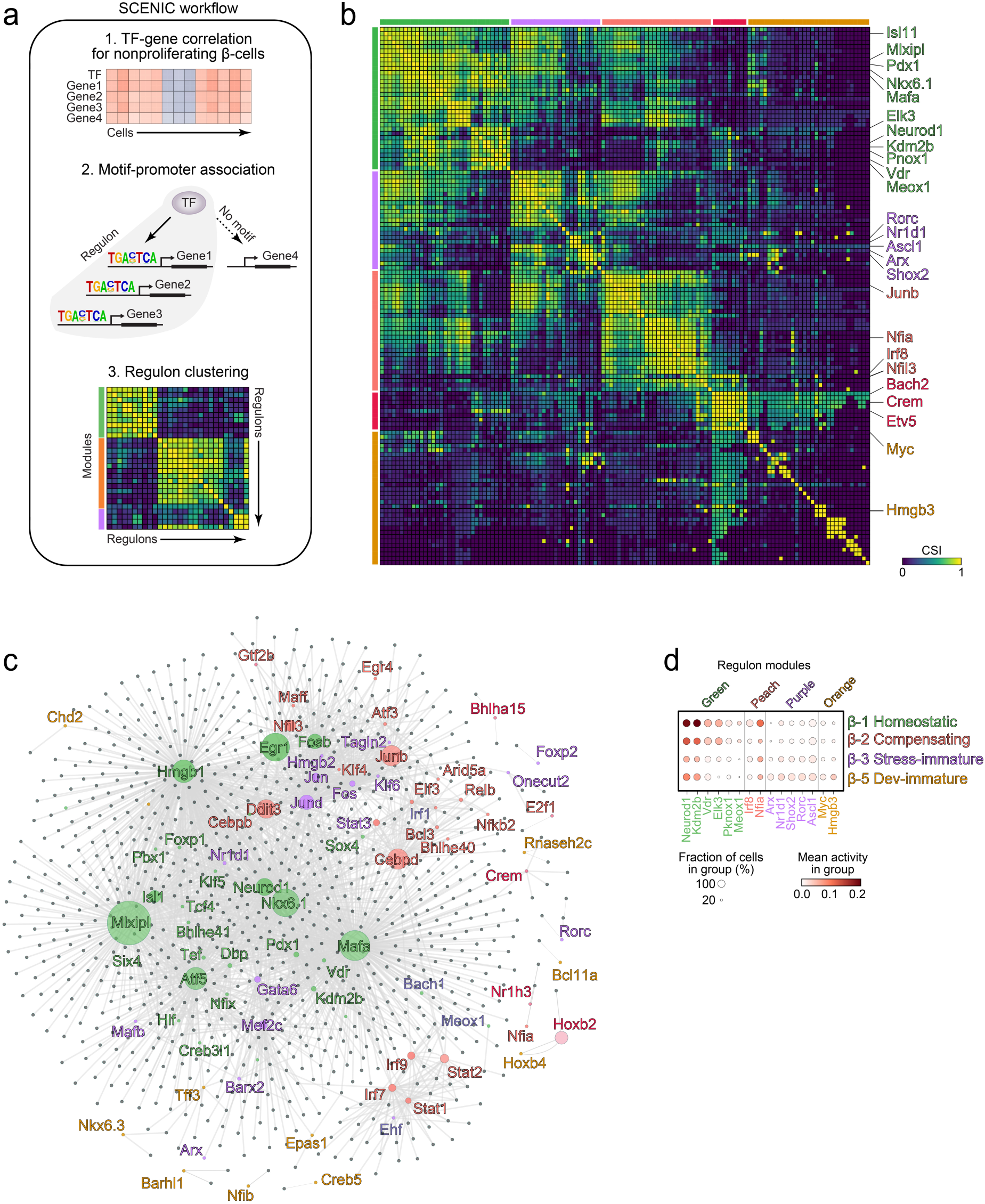
β-cell states exhibit differences in gene regulatory network activities. **(a)** Schematic of TF regulon activity predictions and clustering based on correlated regulon activities among cells. SCENIC, Single-Cell rEgulatory Network Inference and Clustering. **(b)** Correlation plot of all identified regulons clustered into modules. Selected regulons for each module are labeled. Full labeling is shown in Extended Data Fig. 6. CSI, connection specificity index. **(c)** Network visualization of the top 1% of regulons based on SCENIC importance metric. Regulons are colored by module from (b) and target genes are shown in gray. Node sizes indicate Betweenness Centrality. **(d)** Dot plot showing activities for the indicated regulons selected from (b) for each β-cell state.

## Discussion

Through monitoring and manipulating β-cell responses to altered insulin requirements, we have defined β-cell states regulated by environmental workload signals and determined the role of state transitions in diabetes pathogenesis. We found that short- and long-term increases to β-cell workload cause β-cells to transition to a state defined by intermediate expression of mRNAs characteristic of either ‘homeostatic’ or ‘stressed’ β-cells. Experimental manipulation of β-cell states via deletion of *Lsd1* revealed eventual maladaptive effects of β-cell state transitions in response to increased workload. Overall, this study defines the stepwise process of β-cell failure in response to unsustainable workload.

We have identified epigenomic regulation as a key mechanism underlying cell state transitions during changing workloads. Deletion of the chromatin-modifying enzyme *Lsd1* in β-cells mimics the effect of increased workload on β-cell states. The additive effect of *Lsd1* inactivation and insulin resistance in promoting the accumulation of ‘compensating’ and ‘stressed’ β-cells indicates that Lsd1 restrains cell state transitions caused by increased workload. Taken together with our prior findings that Lsd1 dampens the transcriptional response to feeding^20^, this study indicates that Lsd1 opposes the transition between ‘homeostatic’ and ‘compensating’ β-cell states during both insulin resistance and with fasting-feeding rhythms. These results are in agreement with islet single cell multi-ome profiles of H3K27ac or monomethylation of H3 Lys4 (H3K4me1) together with mRNA in mice rendered obese by high-fat diet feeding^38^. Namely, β-cell pseudotime trajectory during obesity corresponds to upregulation of cellular stress markers such as *Fos* and *Hspa5* concomitant with gains in H3K4me1 and H3K27ac at cognate regulatory elements of these genes. As Lsd1 represses transcription of these stress-response genes in β-cells^20^ and facilitates removal of both H3K4me1 and H3K27ac^39,40^, future studies should address how Lsd1-regulated chromatin modifications connect environmental workload signals to β-cell state transitions.

Our findings show that the same mechanisms responsible for β-cell adaptation can become maladaptive when chronically activated. While deletion of *Lsd1* in β-cells of lean, insulin-sensitive mice results in excessive insulin secretion^20^, *Lsd1* inactivation during insulin resistance leads to relative insulin deficiency. Together with analysis of β-cell states, this observation builds the model that ‘compensating’ β-cells initially hypersecrete insulin but eventually fail. Models of constitutive β-cell depolarization such as K_ATP_ channel inactivation^41,42^ or ex vivo tolbutamide treatment^43^ have similarly revealed maladaptive effects of sustained β-cell stimulation. Overall, these findings provide a plausible mechanistic explanation for the enrichment of LSD1-bound sites in human islets for T2D GWAS variants^20^. Namely, genetic variants predisposing to T2D could alter LSD1-dependent interpretation of β-cell workload signals during insulin resistance, leading to shifts in β-cell state that accelerate decompensation.

Through these studies we have generated a mouse islet atlas that should be a valuable resource for understanding how β-cells respond to the pressure to secrete insulin. While previous islet atlases have focused on disease states^9,26,44–46^, this atlas provides further insight into normal physiology as well as the early stages of diabetes pathogenesis. Analysis of models where β-cells successfully adapt to increased workload revealed an intermediate β-cell state that is distinct from both ‘homeostatic’ β-cells that are most abundant in unchallenged mice and from stressed, dysfunctional β-cells that were the focus of prior studies^25,26,47^. While a putative transitional β-cell state was previously inferred from pseudotime trajectory analysis of preclinical diabetes models, these β-cells were exceedingly rare^26^. By contrast, we show that in several models of increased β-cell workload the majority of β-cells assume the ‘compensating’ state. As metabolic decompensation corresponds to accumulation of ‘stressed’ β-cells, the success or failure of β-cell adaptation to increased workload depends on whether the ‘compensating’ β-cell state is sustained. Overall, this atlas provides a foundation for follow-up studies incorporating additional single cell modalities such as proteomic, metabolic, and functional readouts to build a comprehensive understanding of β-cell adaptation and failure in T2D.

## Methods

### Animal studies

All mice used were of mixed strain backgrounds of approximately 50% Bl6N and 50% CD1 for S961 or fasting/feeding studies and 50% C57BLKS/J, 25% C57BL/6N and 25% CD1 for *db/db* studies. The following strains were used in this study: *Lsd1^flox^*^48^*, Pdx1-creER*^49^, and *db/db* (Jackson labs strain 000642). To delete *Lsd1* in *Lsd1^flox/flox^*; *Pdx1-CreER* or *Lsd1^flox/flox^*; *db/db*; *Pdx1-CreER* mice, tamoxifen (Sigma) was injected subcutaneously with 4 doses of 4 mg or 0.15 mg/g body weight, every other day in 10- to 14-week-old mice and 4-week-old mice, respectively. Tamoxifen was dissolved in corn oil at 20 mg/mL for injections. For S961 experiments, mice were implanted with microosmotic pumps (Alzet Model 2002) loaded with S961 peptide or DPBS vehicle to infuse at a rate of 20 nmol/week starting 2 days after the final tamoxifen injection. Blood glucose was monitored with a Bayer Contour glucometer. Fasting and re-feeding was performed as previously described^20^. Briefly, animals were first entrained for 6 days with 12 hours of ad libitum food availability during the dark phase and 12 hours of fasting during the light phase. After entrainment, food was provided at the onset of the dark phase (‘fed’ group) or animals continued to be fasted (‘fast’ group), then islets were isolated 4 hours later at Zeitgeber time (ZT) 16. Only male mice were used for S961 studies and both sexes were used for in vivo *db/db* studies. Sex and age of mice used for in-house-generated and external scRNA-seq datasets for the mouse islet atlas are provided in **Supplemental Table 2**. All animal experiments were approved by the Institutional Animal Care and Use Committee of the University of California, San Diego.

### Metabolic studies

For intraperitoneal glucose tolerance tests, mice were fasted for 6 hours at the onset of the light phase then injected with 1 g glucose per kg body weight. For insulin tolerance tests, mice were injected with 2 U/kg of bovine insulin (Sigma) after 4 hours of fasting. Blood glucose was measured as above at the indicated time points following injections. For serum insulin measurements, blood was collected from the tail vein both prior to and after injection with 1 g glucose per kg body weight following 6 hours of fasting. Serum was prepared by centrifugation and assayed for insulin content by ELISA (ALPCO).

### Islet studies

Islets were isolated and static insulin secretion assays were performed as previously described^20^. Insulin secretion assays were conducted immediately following islet isolations unless otherwise indicated. Secreted insulin was quantified by ELISA (ALPCO).

### Histological studies

Pancreata were harvested and immunostaining was performed and pancreatic insulin content and β-cell mass quantified as previously described^20^. Lsd1 was detected using an anti-Lsd1 antibody (ab17721, Abcam) diluted to 1:1000.

### scRNA-seq sample preparation

For in-house generated scRNA-seq data, islets were isolated essentially as described^20^ with HBSS modified to include 10% FBS and 8.3 mM glucose and to exclude bovine serum albumin. Freshly-isolated islets from the indicated mice (**Supplemental Table 2**) were dissociated using Accutase (StemPro) by incubation at 37**°**C for 10 min with gentle trituration every 2.5 min. Digestion was quenched using DPBS supplemented with 10% FBS, 1 mM EDTA, 25 mM HEPES, and 8.3 mM glucose. Dissociated cells were pelleted then resuspended in quenching buffer containing 1 μM SYTOX Blue (Life Technologies). Cells were sorted by FACS on a BD Influx (BD Biosciences) to exclude dead cells and doublets. Live islet cells were pelleted, resuspended in quenching buffer, then prepared using Chromium Single Cell 3ʹ v3 reagents (10x Genomics) and Chromium X Controller (10x Genomics) according to the manufacturer’s instructions.

### scRNA-seq data processing

#### Data preprocessing

Gene expression count matrices were generated for each of the experimental samples using analysis pipelines and reference genomes provided by 10x Genomics. For islets isolated from S961- or vehicle-treated mice or *db/db* mice with and without β-cell *Lsd1* inactivation the Cell Ranger analysis pipeline v3.0.2 was used for mapping to the mm10 reference 3.0.0 genome. The Cell Ranger analysis pipeline v4.0.0 and the mm10 2020-A genome build was used for the islet single-cell atlas. All subsequent analysis was performed in R (v4.0.2) unless otherwise indicated. The filtered gene expression matrices, which excluded barcodes corresponding to background noise, were processed to remove ambient mRNA contamination using SoupX (v1.5.2)^50^. The contamination fraction for individual samples was estimated based on the expression of hormone and TF markers of the islet endocrine celltypes: α-cells (*Gcg*, *Arx*, *Ttr*), β-cells (*Ins1*, *Ins2*, *Iapp*), δ-cells (*Sst*) and PP-cells (*Ppy*, *Pyy*). The counts were then adjusted with ‘adjustCounts()’ function. Following ambient mRNA correction, we filtered out doublets by applying Scrublet (v0.2.1)^51^ on each individual sample. The ‘expected_doublet_rate’ parameter was manually adjusted based on the total number of filtered cells obtained from Cell Ranger.

#### Data normalization and integration

Sample integration and analysis was performed using Seurat (v4.1.0)^52^. The ambient mRNA and doublet filtered matrices of individual samples were imported as Seurat objects with the ‘CreateSeuratObject()’ function, while filtering out: (i) features detected in less than 10 cells, (ii) cells with less than 500 features detected, and (iii) outlier cells based on UMI counts, feature counts and with >15% of counts associated with the mitochondrial genome (mt-genes). Next, the cell-x-gene count matrices were log-normalized using the ‘NormalizeData()’ function and scaled across the top 3000 highly variable genes (HVGs) using the ‘ScaleData()’ function. For sample integration, anchors were obtained using the ‘FindIntegrationAnchors()’ function using a reciprocal PCA (rpca) approach, and were subsequently integrated using the ‘IntegrateData()’ function.

The integrated data was further scaled across the HVGs, following which principal component analysis (PCA) was performed using the ‘RunPCA()’ function. The top 30 principal components (PCs) were used to build a neighborhood graph with the ‘FindNeighbors()’ function. The computed graph was then used to reduce the dimensionality of the integrated data from 30 PCs to a two-dimensional projection via the Uniform Manifold and Approximation Projection (UMAP) algorithm^53^, implemented in the ‘RunUMAP()’ function. Cell clusters were obtained with the ‘FindClusters()’ function using a resolution of 0.2.

#### β-cell state analysis

β-cells were identified based on the expression of *Ins1/2* exclusive of other islet endocrine hormone markers (*Gcg, Sst*, and *Ppy*). The remaining β-cells were re-integrated by identifying β-cell specific HVGs, then PCA was performed and a β-cell specific UMAP embedding was computed using the top 20 PCs. β-cells were clustered using the ‘FindClusters()’ function at a resolution of 0.1 and state-specific markers were defined by Wilcoxon rank sum test using the ‘FindAllMarkers()’ function.

#### Graph abstraction and pseudotime

The β-cell-specific Seurat objects from non-proliferating cell clusters were converted to anndata (v0.7.8) H5AD objects using SeuratDisk (v0.0.0.9021)^54^. The β-cell H5AD objects were then imported using Scanpy (v1.7.2)^55^ for downstream analysis. To analyse the connectivity between the identified β-cell states, a k-nearest neighbor graph was re-computed with ‘sc.pp.neighbors’. Based on this underlying neighbor graph, Partition-based Graph Abstraction (PAGA)^27^ was performed using ‘sc.tl.paga’ to estimate the connectivity between the β-cell states, thereby generating a simplified representation of the transition between cell states.

For pseudotime analysis, a root cell was defined within the β-1 homeostatic state, and diffusion pseudotime (DPT)^56^ was computed using ‘sc.tl.dpt’ with 15 diffusion components (‘n_dcs = 15’). Using the DPT, cells were ordered along a trajectory based on diffusion distances on the neighbor graph and scaled gene expression was visualized using ‘sc.pl.paga_path’.

#### Gene regulatory network inference and regulon module analysis

pySCENIC (v0.11.2) was used to infer GRNs for the non-proliferating β-cells^57^. Regulons (TF-target genes) were generated then their activities in individual cells were computed and binarized (assigned ON or OFF value, per regulon per cell). The regulon modules were identified based on connection specificity index (CSI)^58^, which were computed from the Pearson correlation coefficient (PCC) of the activity scores for each pair of regulons, upon which hierarchical clustering with complete linkage was performed.

A heatmap depicting the CSI of the five regulon modules across the non-proliferating β-cells was generated with ‘ComplexHeatmap’ R package (v2.6.2)^59^. The network plot was generated with Cytoscape (v3.10.0)^60^ from the top 1% of the TF-target associations from the co-expression modules based on the reported Importance Metric as described elsewhere^61^. The betweenness centrality of the nodes in the network were computed within Cytoscape with the Network Analyzer module. The mean activity of regulons across β-cell states or samples were computed using ‘aggregate()’ function in base R and dot plots depicting the mean regulon activity were generated with custom plotting scripts.

## Gene ontology

Enrichment of pathways (KEGG and Reactome) and gene ontology terms (Gene Ontology - Biological Process) for markers overexpressed in each β-cell state was determined using default parameters in Metascape (v3.5)^62^.

**Supplementary Information** will be linked to the online version of the paper upon publication.

## Acknowledgements

We thank members of the Sander and Wortham labs for helpful discussions and Kristen Wells for comments on the manuscript. We acknowledge support of the UCSD IGM Genomic Center (P30 DK064391) for scRNA-seq. This work was supported by a Diabetes Research Center Pilot and Feasibility grant P30 DK063491 (M.W.), JDRF grant SRA-2021-1052-S-B (M.S. and M.W.), NIH grants DK068471 (M.S. and M.W.) and DK078803 (M.S.). This publication includes data generated at the UCSD IGM Genomics Center utilizing an Illumina NovaSeq 6000 that was purchased with funding from an NIH SIG grant (S10 OD026929).

## Author Contributions

S.S., M.S., and M.W. conceived the study, were responsible for its overall design, and prepared the manuscript. F.L., A.R.H., and M.W. performed mouse experiments. F.L. performed islet isolations. A.R.H., D.P.P., H.Z., I.O., D.P.P., and M.W. performed islet experiments and hormone measurements. S.S., M.M., Y.S., M.W. performed bioinformatics analysis.

## DATA AVAILABILITY

scRNA-seq data will be deposited in GEO prior to publication.

