## Extended Data Figures for "Workload-induced changes to cell state contribute to β-cell failure in diabetes"

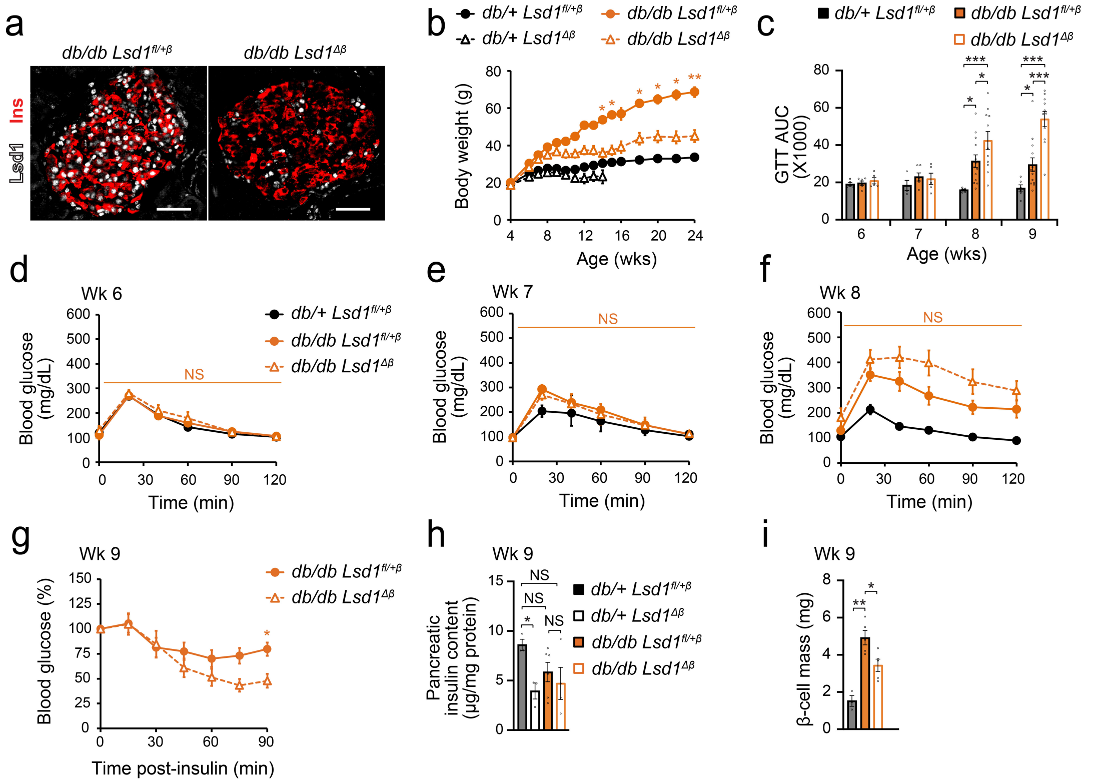


**Extended Data Figure 1. *Lsd1* inactivation in β-cells accelerates the onset of glucose intolerance in *db/db* mice. Related to Figure 1.**

**(a)** Immunofluorescence staining for insulin (Ins) and Lsd1 on pancreas sections from TM-treated *$Lsd1^{fl/+}$*; *db/db*; *Pdx1-CreER* (*db/db $Lsd1^{fl/+\beta}$*) mice and TM-treated *$Lsd1^{fl/fl}$*; *db/db*; *Pdx1-CreER* (*db/db $Lsd1^{\Delta\beta}$*) mice two days after TM treatment as in Fig. 1a. Scale bar, 50 µm. TM, tamoxifen. Representative image from $n = 1$ mouse per group.

**(b)** Longitudinal body weight measurements of *db/+ $Lsd1^{fl/+\beta}$* ($n = 7$), *db/+ $Lsd1^{\Delta\beta}$* ($n = 14$) *db/db $Lsd1^{fl/+\beta}$* ($n = 8$), and *db/db $Lsd1^{\Delta\beta}$* ($n = 11$). Wks, weeks. *$p<0.05$, **$p<0.01$ between *db/db $Lsd1^{fl/+\beta}$* and *db/db $Lsd1^{\Delta\beta}$* mice.

**(c - f)** Longitudinal glucose tolerance tests in mice of the indicated genotypes shown as (c) area under the curve (AUC) for all time points or (d - f) blood glucose levels following intraperitoneal glucose injection for each time point. AUC, area under the curve. *db/+ $Lsd1^{fl/+\beta}$* wk 7: $n = 4$ mice, *db/db $Lsd1^{fl/+\beta}$* wks 6 and 7: $n = 6$ mice, *db/db $Lsd1^{\Delta\beta}$* wk 8: $n = 11$ mice, *db/db $Lsd1^{fl/+\beta}$* wks 8 and 9: $n = 13$ mice, *db/+ $Lsd1^{fl/+\beta}$* wk 9: $n = 6$ mice, all other groups: $n = 5$ mice. NS, not significant between *db/db $Lsd1^{fl/+\beta}$* and *db/db $Lsd1^{\Delta\beta}$* mice.

**(g)** Insulin tolerance tests in 9-week-old *db/db $Lsd1^{fl/+\beta}$* ($n = 13$), and *db/db $Lsd1^{\Delta\beta}$* ($n = 11$) mice. Blood glucose levels are shown relative to the 0 min time point. *$p<0.05$ between *db/db $Lsd1^{fl/+\beta}$* and *db/db $Lsd1^{\Delta\beta}$* mice.

**(h and i)** Pancreatic insulin content (h) and β-cell mass (i) in 9-week-old mice of the indicated genotypes. *db/db $Lsd1^{fl/+\beta}$*: $n = 6$ (h) or $n = 5$ (i) mice, *db/db $Lsd1^{\Delta\beta}$* $n = 4$ (h) or $n = 5$ (i) mice, all other groups: $n = 3$ mice.

Significance was determined by one-way ANOVA followed by Student's t-test with Welch's correction for unequal variance as necessary followed by Dunnett's multiple comparisons test (c, h, i) or two-way ANOVA for genotype interaction with time followed by Sidak's multiple comparisons test (b, d - g).

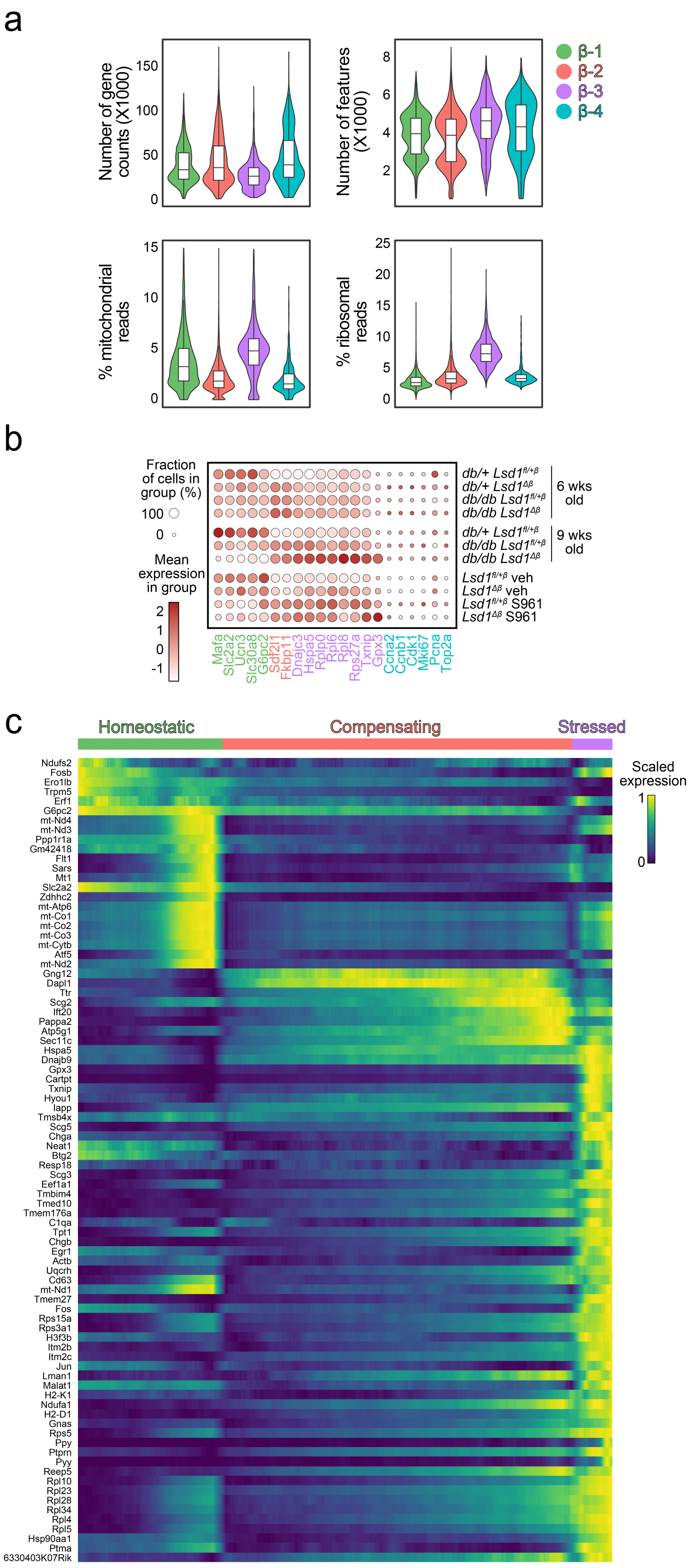


**Extended Data Figure 2. Validation of β-cell state classifications. Related to Figure 2.**

**(a**) Violin plots of quality control metrics for each β-cell state from Fig. 2b.
**(b)** Dot plot showing expression of mRNAs from Fig. 2d for each sample. Gene symbol colors indicate the state enriched for each corresponding mRNA.
**(c)** mRNA expression changes along the diffusion pseudotime trajectory for the three most abundant β-cell states in Fig. 2b. Cells were ordered as in [27].

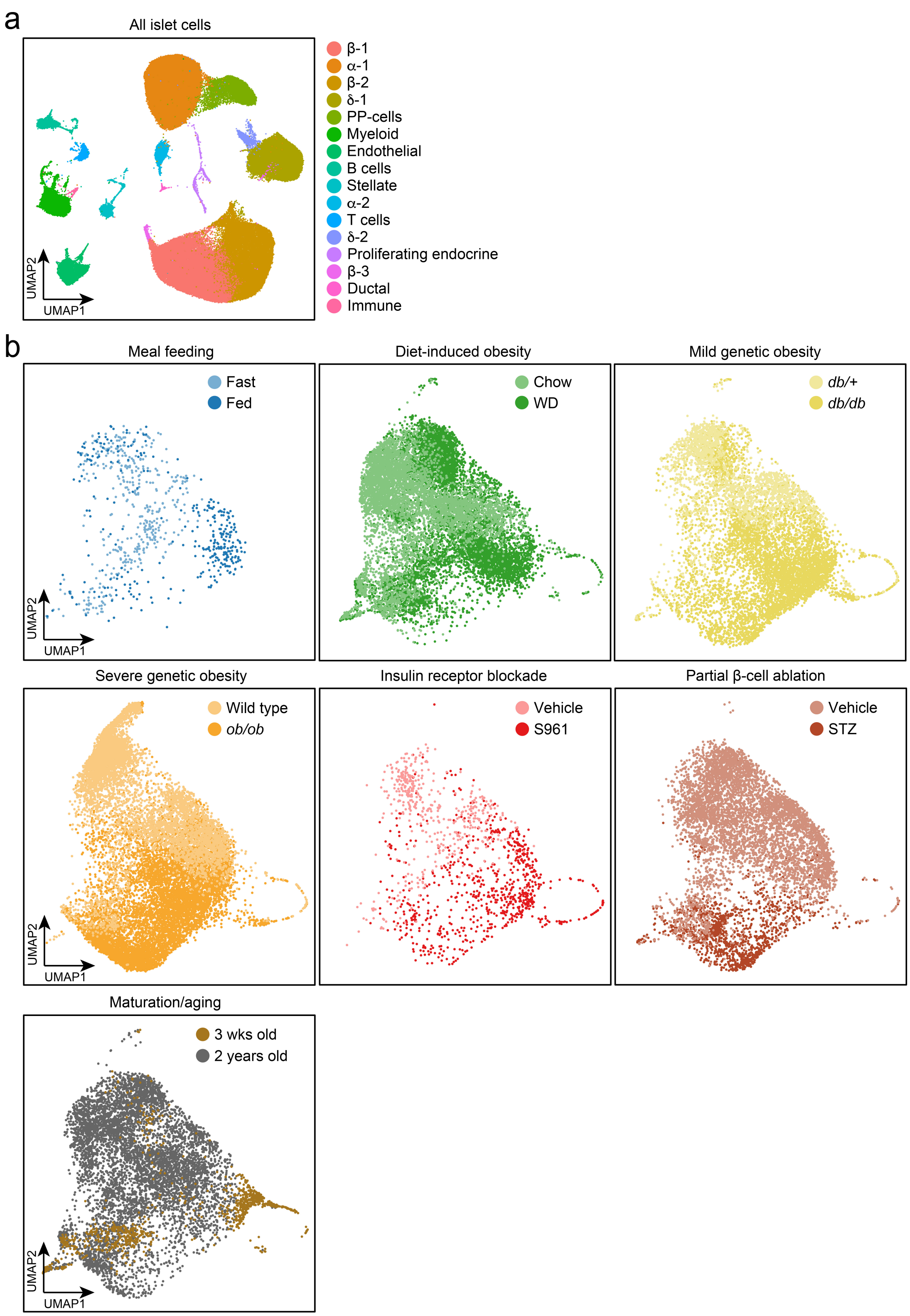


**Extended Data Figure 3. Validation of data integration. Related to Figure 3.**

**(a)** UMAP of integrated and clustered islet cells colored by cell type and state. UMAP, uniform manifold approximation and projection.
**(b)** UMAPs of re-integrated β-cells from the indicated datasets colored by genotype or treatment group.

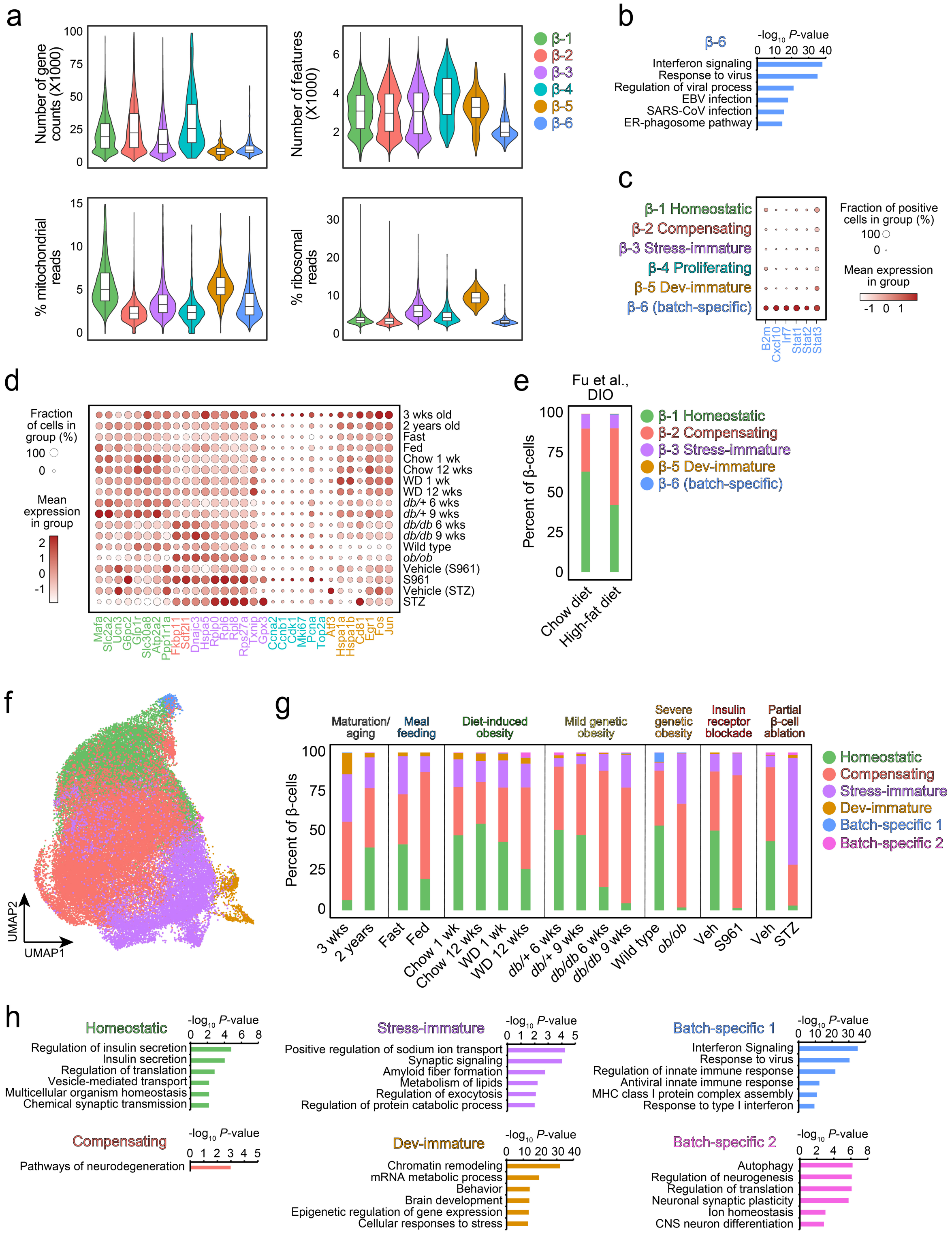


**Extended Data Figure 4. Validation of β-cell state classifications in the mouse islet atlas. Related to Figure 3.**

(**a**) Violin plots of quality control metrics for each β-cell state from Fig. 3b.

**(b)** Enriched gene ontologies and pathways for mRNAs more highly expressed in the β-6 subset relative to all other β-cells.
**(c)** Dot plot showing expression of representative mRNAs from gene ontologies and pathways selected from (b).
**(d)** Dot plot showing expression of mRNAs from Fig. 3d for each sample. Gene symbol colors indicate the state enriched for each corresponding mRNA.
**(e)** Abundance of each β-cell subset as a proportion of total β-cells for β-cells from chow- or high-fat diet-fed mice [63] mapped to the integrated dataset in Fig. 3f. DIO, diet-induced obesity.
**(f and g)** UMAP of clustered nonproliferating β-cells colored by state after regressing mitochondrially encoded mRNAs and ribosomal protein genes (f) and abundance of each β-cell state as a proportion of total β-cells for the indicated treatments and genotypes (g). UMAP, uniform manifold approximation and projection.
**(h)** Enriched gene ontologies and pathways for mRNAs more highly expressed in each state relative to all other β-cells. The top six most significantly enriched categories are shown.

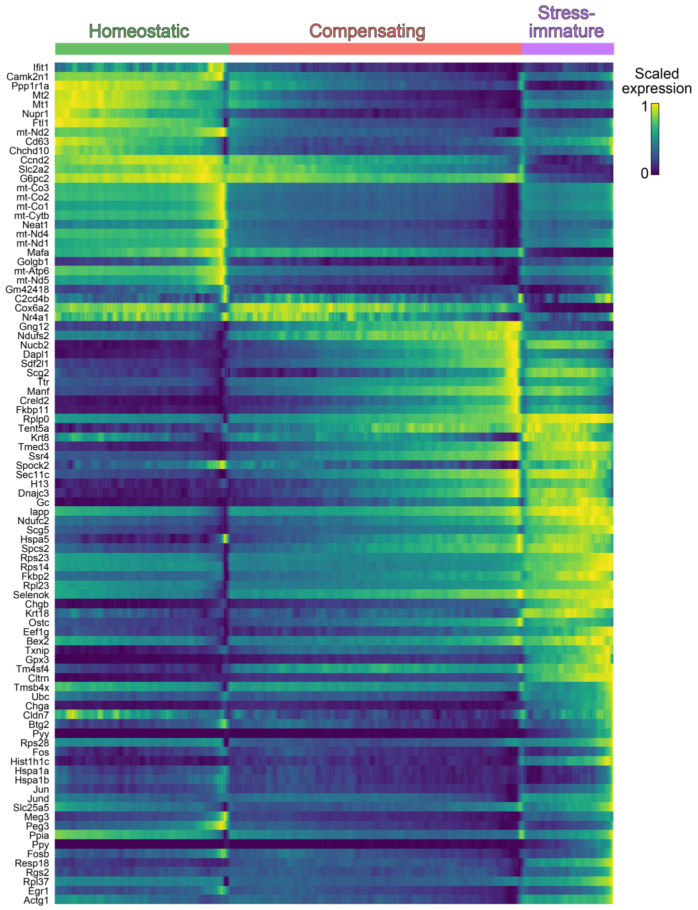


**Extended Data Figure 5. Diffusion pseudotime trajectory of β-cells in the mouse islet atlas. Related to Figure 3.**

mRNA expression changes along the diffusion pseudotime trajectory for the three most abundant β-cell states in the mouse islet atlas (β-1 through β-3). Cells were ordered as in [27].

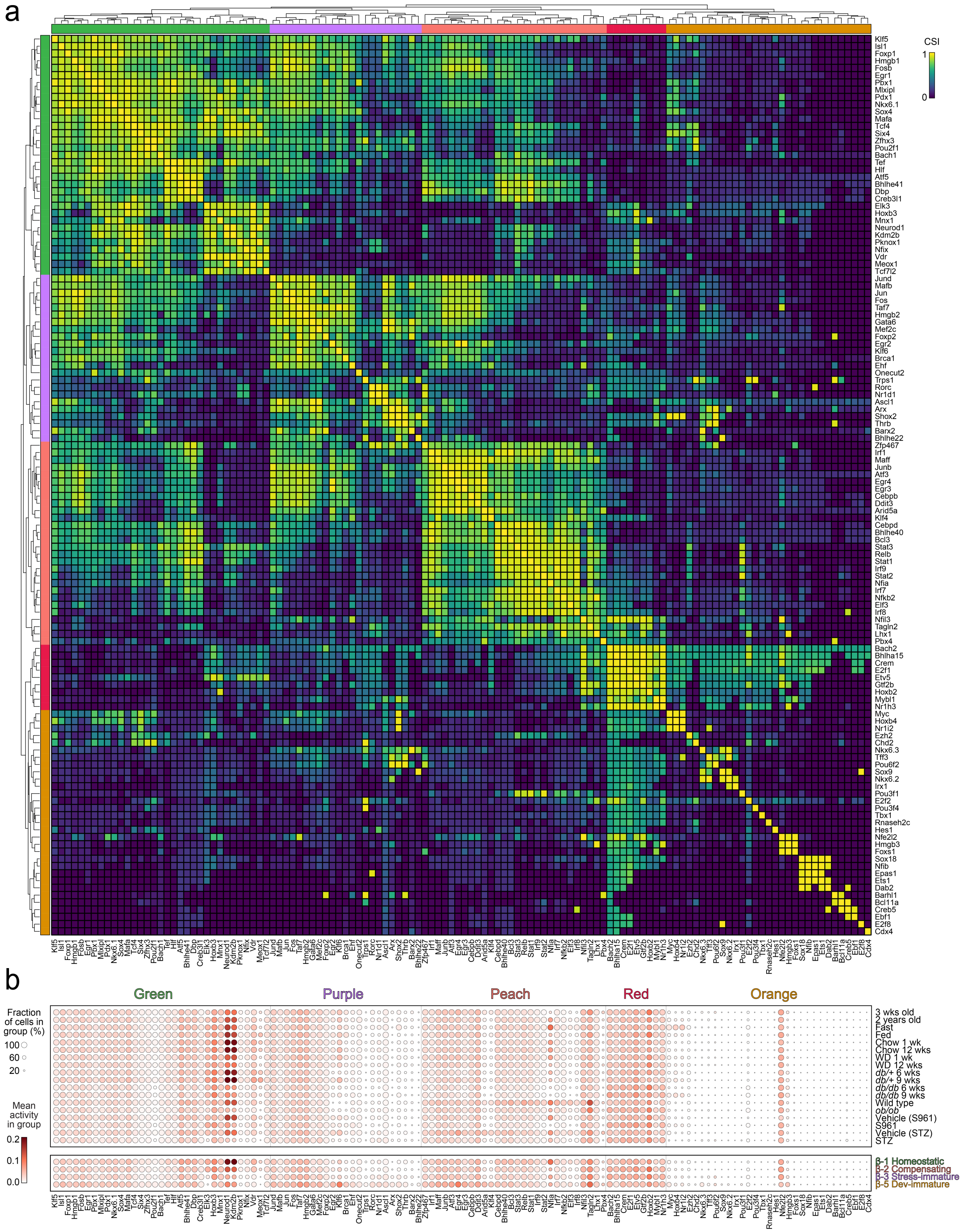

**Extended Data Figure 6. Transcription factor activities correspond to distinct β-cell states. Related to Figure 4.**

**(a)** Correlation plot of pairwise regulon activities among nonproliferating β-cells from experimental groups in Fig. 3a clustered into co-expression modules using CSI (connection specificity index).
**(b)** Dot plot showing showing activities for the indicated regulons from (a) in nonproliferating β-cells from the indicated experimental groups and states.
